# Structural basis for differential inhibition of THIK1 by chemically related inhibitors

**DOI:** 10.64898/2026.09.21.753137

**Authors:** Ran Zhang, Xiangyun Fang, Ruiheng Zhang, Zixue Wang, Haichao Jin, Boxuan Wang, Wei Wang, Xiaofang Zhong, Jin Wang, Baobin Li

**Affiliations:** Huadong Hospital, Institute for Translational Brain Research, State Key Laboratory of Medical Neurobiology, MOE Frontiers Center for Brain Science, Fudan University, Shanghai, China; School of Science, China Pharmaceutical University, Nanjing, China; State Key Laboratory of Crop Stress Adaptation and Improvement, the Zhongzhou Laboratory for Integrative Biology, School of Life Sciences, Henan University, Kaifeng, China; College of Chemistry and Molecular Engineering, Peking University, Beijing, China

## Abstract

TWIK-related halothane-inhibited potassium channel 1 (THIK1) is a two-pore domain potassium (K2P) channel highly expressed in microglia that contributes to microglial homeostasis and neuroinflammatory responses. Pharmacological inhibition of THIK1 suppresses tonic K^+^ currents and attenuates NLRP3-dependent IL-1β release, highlighting THIK1 as a potential therapeutic target for neuroinflammatory disorders. However, the structural basis of THIK1 inhibition remains poorly understood. Here we report cryo-electron microscopy (cryo-EM) structures of full-length human THIK1 in complex with C101248, a commercially available preclinical compound, and CVN293, an inhibitor currently advancing toward Phase II clinical trials. The two inhibitors occupy a similar site within the inner vestibule formed by TM2 and TM4 beneath the selectivity filter, implicating this region in THIK1 inhibition. Consistent with their closely related chemical scaffolds and shared binding site, both inhibitors remodel the Y273 inner gate; however, they differ in their effects on the C-terminal gate. Electrophysiological analyses further demonstrate that the C-terminal gate contributes differently to inhibition by the two compounds. Together, our findings provide structural and functional insights into how chemically related inhibitors can exert different effects on THIK1 despite engaging a common binding site.

## Introduction

THIK1 (encoded by KCNK13) is a member of the two-pore potassium ion channel family, which is specially expressed in microglia (1, 2). This channel provides the major tonic K⁺ conductance of microglia, thereby setting the resting membrane potential (2). THIK1 plays an important role in regulating the functional state of microglia (3–5). Under physiological conditions, microglia promote a neuroprotective phenotype, whereas under pathological conditions, they trigger inflammatory responses (6, 7). Recent studies have implicated THIK1 in neuroimmune disease, with elevated expression observed in Alzheimer’s disease (AD) (8), indicating that THIK1 may contribute to disease-associated microglial activation. Consequently, selective blockade of microglial THIK1 at early stages of PD may mitigate inflammatory responses, slow disease progression and provide therapeutic benefit (2, 9). Compared with other microglial regulators, including the purinergic receptors P2Y12 and P2X7 (10, 11), THIK1 represents a more promising therapeutic target owing to its restricted expression in microglia, whereas P2Y12 is also expressed in platelets (12) and P2X7 is broadly expressed across multiple immune cell types (13).

Given the growing therapeutic relevance of THIK1, substantial efforts have been made to identify potent inhibitors targeting THIK1. C101248, the first reported selective THIK1 inhibitor, suppresses THIK1-mediated currents and attenuates NLRP3-dependent interleukin-1β release from microglia, but it was developed primarily as a tool compound rather than a clinical candidate (9). Subsequent optimization of this scaffold led to CVN293, a potent, selective and brain-penetrant clinical candidate with improved drug-like properties. In Phase I studies, CVN293 was well tolerated in healthy adults and exhibited measurable cerebrospinal fluid (CSF) penetration (14, 15). Collectively, THIK1 is a tractable target for modulating microglial activation and neuroinflammatory signaling.

Understanding the molecular mechanisms of THIK1 inhibition is important for the development of potent and selective THIK1 inhibitors. Previously, we determined the structures of full-length human THIK1 (hTHIK1) in both open and closed conformations, providing a structural framework for THIK1 gating (16). Here, we report cryo-electron microscopy (cryo-EM) structures of hTHIK1 in complex with the chemically related inhibitors C101248 and CVN293 at overall resolutions of 2.99 Å and 2.81 Å, respectively (Table 1). Our structural and functional analyses reveal how these chemically related inhibitors engage the inner vestibule and differentially modulate THIK1 conformation, providing structural insights into the molecular mechanism of THIK1 inhibition.

**Table 1.**
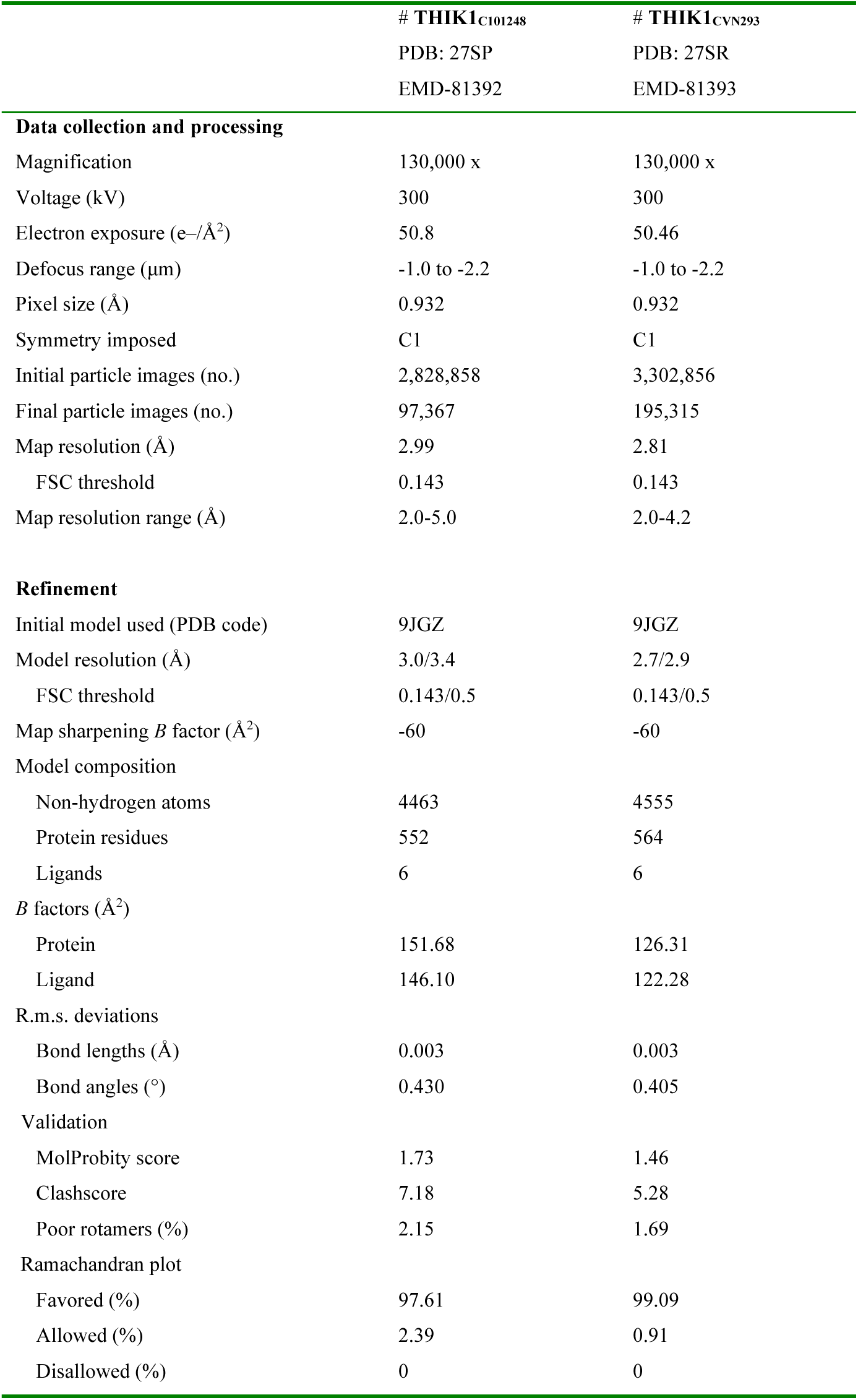
Cryo-EM data collection, refinement and validation statistics.

## Results

### C101248 recognition and inhibition of THIK1

C101248 was identified as the first potent and selective small-molecule inhibitor of THIK1. It blocks both tonic and ATP-evoked K⁺ currents in a concentration-dependent manner and exhibits rapid association with and slow dissociation from THIK1 (9). To further characterize its inhibitory properties, we performed whole-cell patch-clamp electrophysiology in HEK293 cells expressing full-length hTHIK1. Whole-cell patch-clamp recordings in HEK293 cells gave IC₅₀ values of 348.8 nM for C101248 and 427.6 nM for CVN293 (Fig. 1A and 2C). Both values are higher than those reported in thallium-flux assays (9, 14), reflecting differences between electrophysiological and flux-based measurements. Selectivity test showed that C101248 minimally inhibited constitutive currents mediated by hTREK1 or hTASK2 in HEK293 cells, supporting its selectivity for THIK1 over K2P channels examined (Fig. 1A).

**Fig. 1.**
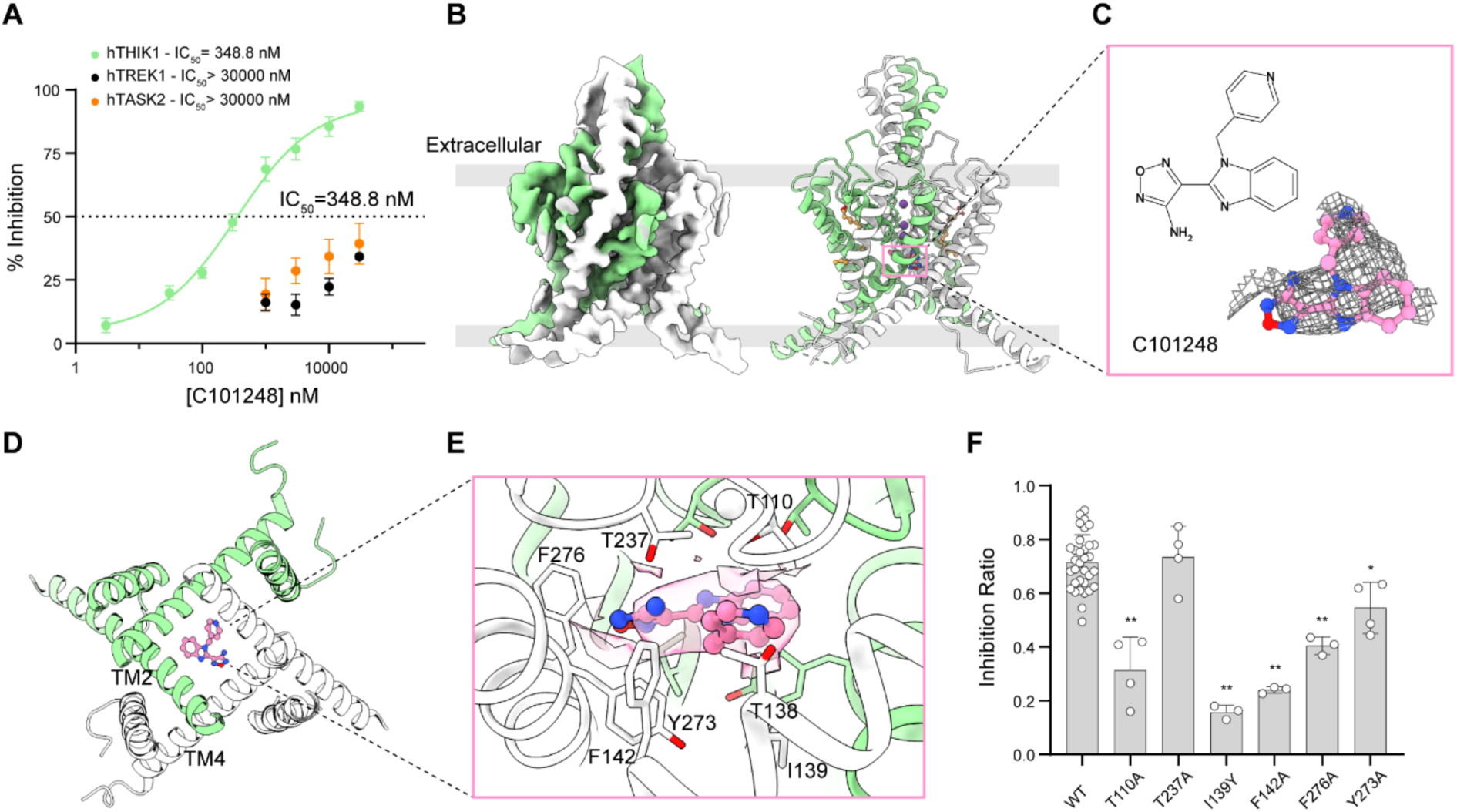
Structural and functional characterization of C101248 binding to THIK1. **(A)** Inhibitory effects of C101248 on hTHIK1, hTREK1 and hTASK2 currents measured by whole-cell patch-clamp electrophysiology in HEK293 cells. Data are presented as mean ± SD (n = 3-4). **(B)** Cryo-EM density map of THIK1_C101248_ viewed parallel to the membrane plane (left); Overall ribbon model of THIK1_C101248_ with protomers shown in light green and white (right). **(C)** Chemical representation and resolved cryo-EM density of C101248. **(D)** Cross-sectional view of the C101248-binding pocket in THIK1, viewed from the extracellular side. **(E)** Magnified view of the binding pocket, with C101248 (pink) and interacting residues (light green) displayed as sticks. The corresponding cryo-EM density for C101248 is shown as a mesh. **(F)** Comparison of inhibition ratio at 50 mV in THIK1_WT_ and binding-pocket mutants. Inhibition ratio refers to the ratio of the current after application of 1 μM C101248 to the current before application. Data are presented as mean ± SD. The number of independent cells tested for each group is indicated by individual symbols (from left to right: n = 33, 4, 4, 3, 3, 3, 4, respectively). Statistical significance was determined using one-way ANOVA followed by Dunnett’s multiple comparisons test. \**P* < 0.05 and \*\**P* < 0.01 *vs.* WT.

**Fig. 2.**
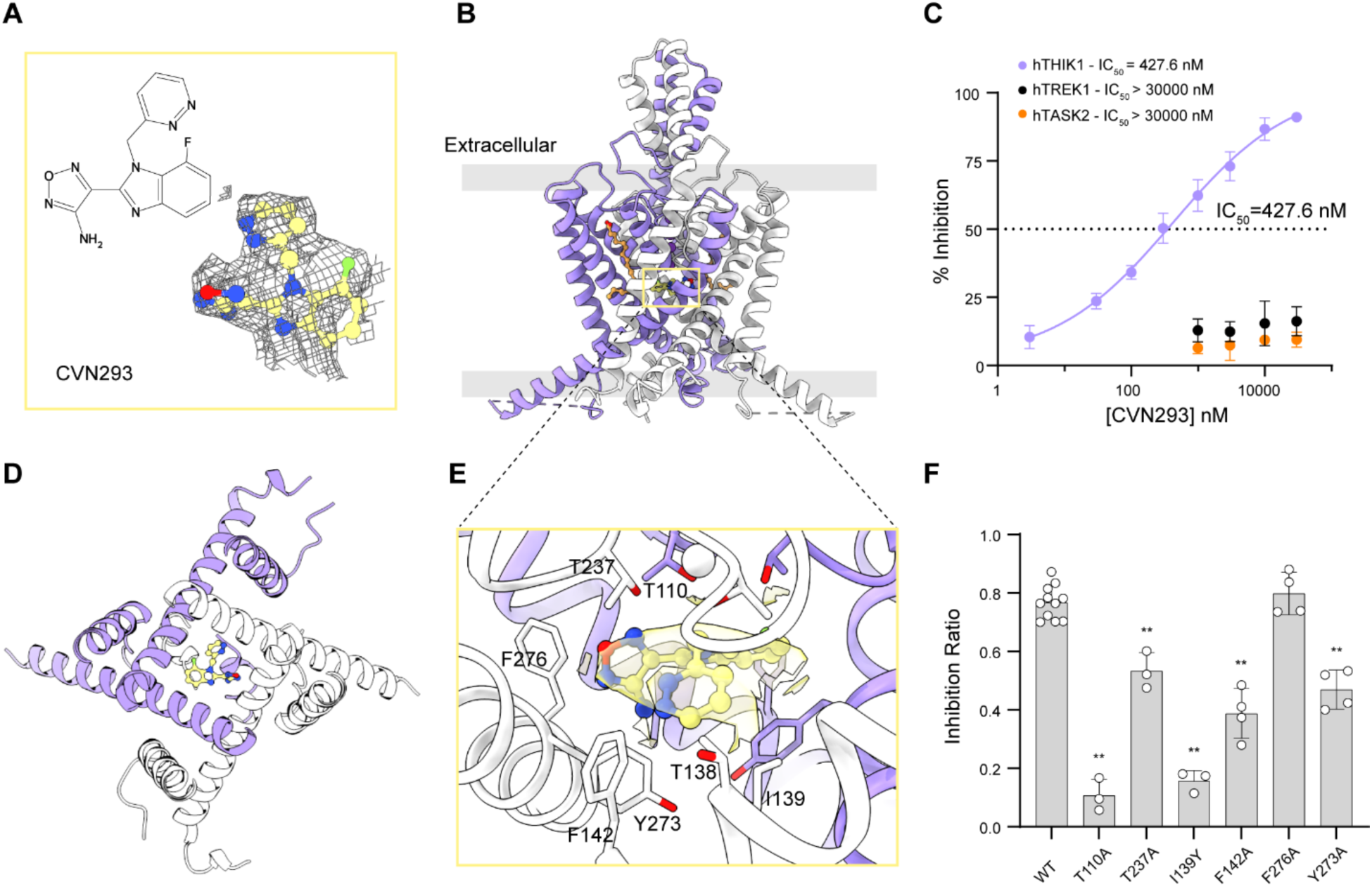
Structural and functional characterization of CVN293 binding to THIK1. **(A)** Chemical representation and resolved cryo-EM density of CVN293. The density is shown as a grey mesh. **(B)** Ribbon model of THIK1_CVN293_ viewed parallel to the membrane plane, depicting the two protomers in purple and white and bound CVN293 in yellow. **(C)** Inhibitory effects of CVN293 on hTHIK1, hTREK1 and hTASK2 currents measured by whole-cell patch-clamp electrophysiology in HEK293 cells. Data are presented as mean ± SD (n = 3-4). **(D)** Cross-sectional view of the CVN293-binding pocket in THIK1, viewed from the extracellular side. **(E)** Close-up view of the binding pocket showing CVN293 (yellow) and interacting residues (purple) in stick representation. Cryo-EM density corresponding to CVN293 is shown as a semi-transparent surface. **(F)** Comparison of inhibition ratio at 50 mV in THIK1_WT_ and binding-pocket mutants. Inhibition ratio refers to the ratio of the current after application of 10 μM CVN293 to the current before application. Data are presented as the mean ± SD, with individual symbols indicating the number of independent cells tested per group (n = 11, 3, 3, 3, 4, 4, 4, from left to right). Statistical analysis was performed using a one-way ANOVA, followed by Dunnett’s post-hoc test. \**P* < 0.05 and \*\**P* < 0.01 denote significant differences compared to the WT group.

As the first potent and selective small-molecule inhibitor of THIK1, C101248 provides an important framework for understanding the structural basis of THIK1 inhibition. We therefore expressed full-length THIK1 in HEK293S GnTI⁻ cells and purified the channel for single-particle cryo-EM analysis, eventually determined the structure of THIK1 in complex with C101248 in LMNG/CHS detergent with a resolution of 2.99 Å (Fig. 1*B*; Table 1; Fig. S1). Given the asymmetry of the ligand and the possibility of symmetry breaking in THIK1, we performed ab-initio reconstruction and non-uniform refinement with C1 symmetry imposed throughout (Fig. S1).

Like other K2P channels, THIK1_C101248_ adopts the canonical homodimeric architecture (Fig. 1*B*), with each protomer containing four transmembrane helices (TMs) (17–21). Inspection of the cryo-EM map, we identified a strong density beneath the selectivity filter (SF) that was readily assigned to C101248 based on its shape (Fig. 1*C*). C101248 binds within the inner vestibule formed primarily by TM2 and TM4, a key regulatory hub in K2P channels that couples intracellular gating to the SF (Fig. 1*D*). Its occupancy is expected to impede K⁺ conduction from the intracellular side, thereby inhibiting channel activity (22). A cluster of polar and hydrophobic residues surround the ligand, including T110, T138, I139, F142, T237, Y273 and F276 (Fig. S2, *A* and *B*). Among them, Y273 in TM4 forms the base of the binding pocket, which is identified as inner gate II (16, 23, 24). The aromatic core of C101248 lies in close proximity to Y273’s side chain allowing for potential π–π stacking, and its amino group forms a hydrogen bond with T237 (subunit B). Additional van der Waals contacts with T138, I139 and F142 further stabilize the ligand within the cavity (Fig. 1*E* and Fig. S2, *B* and *C*). Comparing residues around the inner vestibule of THIK1_C101248_ and THIK1_WT_ (PDB ID: 9JGZ) reveals a subtle outward shift of pocket-lining side chains in TM2, TM4 and beneath SF upon C101248 binding, with Y273 undergoing the most pronounced rearrangement, moving downward by ∼3.5 Å (Fig. S2, *C* and *D*). Consequently, the distance between the hydroxyl groups of Y273 from each subunit increases from 2.1 Å in THIK1_WT_ to 5.0 Å in THIK1_C101248_ (Fig. S2, *D* and *E*). These conformational changes expand the inner vestibule to accommodate the inhibitor and may couple its binding to gating transitions in distal regions of the channel (16, 23, 24).

To assess the contribution of pocket-lining residues to C101248 inhibition, we introduced mutations within the vestibular binding pocket and evaluated their effects on C101248-mediated inhibition. Most mutations reduced C101248-mediated inhibition, supporting the observed binding mode in the THIK1_C101248_ structure (Fig. 1*F*).

### Conserved vestibular recognition of the clinical inhibitor CVN293

Based on C101248, introducing a fluoride at position 7 of the benzimidazole core and optimizing heterocyclic groups containing a second ring nitrogen led to CVN293, another selective THIK1 inhibitor developed by Cerevance, which designed to target NLPR3-mediated neuroinflammation and demonstrates improved metabolic stability and pharmacokinetic properties (14). In a Phase I study, CVN293 was well tolerated in healthy adult subjects and capable of penetrating the CSF, indicating its potential utility for central nervous system therapy (15). To understand how CVN293 inhibits THIK1 at the molecular level and compare the inhibitory mechanisms of structurally related compounds, we determined the structure of THIK1 in complex with CVN293 in LMNG/CHS detergent with a resolution of 2.81 Å (Fig. 2, *A* and *B*; Table 1; Fig. S3). The sample preparation and data processing of THIK1_CVN293_ followed the same protocol as those used for THIK1_C101248_. Whole-cell patch-clamp electrophysiology yielded an IC_50_ of 427.6 nM for CVN293, slightly higher than that of C101248 (Fig. 2*C*). Like C101248, CVN293 exhibited pronounced selectivity for THIK1 over the other K2P channels examined (Fig. 2*C*).

As observed for C101248, CVN293 also occupies the vestibule beneath the SF (Fig. 2*D* and Fig. S4*A*). It is accommodated within a pocket predominantly formed by hydrophobic residues, including I139, F142, Y273 and F276, stabilizing CVN293 through extensive hydrophobic interactions and van der Waals contacts (Fig. S4*B*). The aromatic side chains of F142 and Y273 are positioned near the aromatic ring of CVN293, with the distances ranging from 3.4 to 4.5 Å, consistent with potential π–π interactions (Fig. S4*C*) (25). T110 and T237 on the SF are positioned above CVN293, with their side-chain hydroxyl groups positioned to interact with the ligand (Fig. 2E and Fig. S4, *B* and *C*). Notably, these interacting residues largely overlap with the C101248-binding site and undergo a similar outward rearrangement relative to the wild-type structure. Y273 adopts a similar rearrangement in THIK1_C101248_, accompanied by a marked expansion of the inter-subunit hydroxyl distance (Fig. S4*D*). Single mutations of either selectivity filter residues (T110 and T237) or vestibule-lining residues (I139, F142 and Y273) markedly attenuated CVN293 inhibition (Fig. 2*F* and Fig. S4*E*), supporting a role for these residues in CVN293-mediated inhibition.

Considering the high selectivity of CVN293 and C101248 for THIK1 over other K2P channels, we examined the conservation of residues forming the inhibitor-binding pocket across the K2P family. Sequence alignment revealed that most ligand-contacting residues are largely restricted to the THIK subfamily (Fig. S5*A*). These findings provide a structural basis for the preferential inhibition of THIK channels by C101248 and CVN293. Moreover, the residues forming the ligand-binding pocket are highly conserved across human, mouse, and rat KCNK13, with an overall amino acid identity of ∼85% (Fig. S5*B*). This conservation suggests that the vestibular pocket represents a conserved pharmacological site across species and may facilitate structure-guided development of subtype-selective K2P modulators (26).

### Differential conformational effects of C101248 and CVN293 on THIK1

To further investigate differences in the inhibitory mechanisms of C101248 and CVN293, we superposed the two inhibitor-bound structures with each other and with THIK1_WT_ (Fig. S6). In addition to the previously described downward displacement of Y273 at inner gate II upon inhibitor binding, the two inhibitors exhibited distinct effects on inner gate I. The C-terminal region was largely unresolved in the THIK1_C101248_ structure, whereas the THIK1_CVN293_ structure retained a well-defined closed conformation of inner gate I (Fig. 3, *A* to *D* and Fig. S6).

**Fig. 3.**
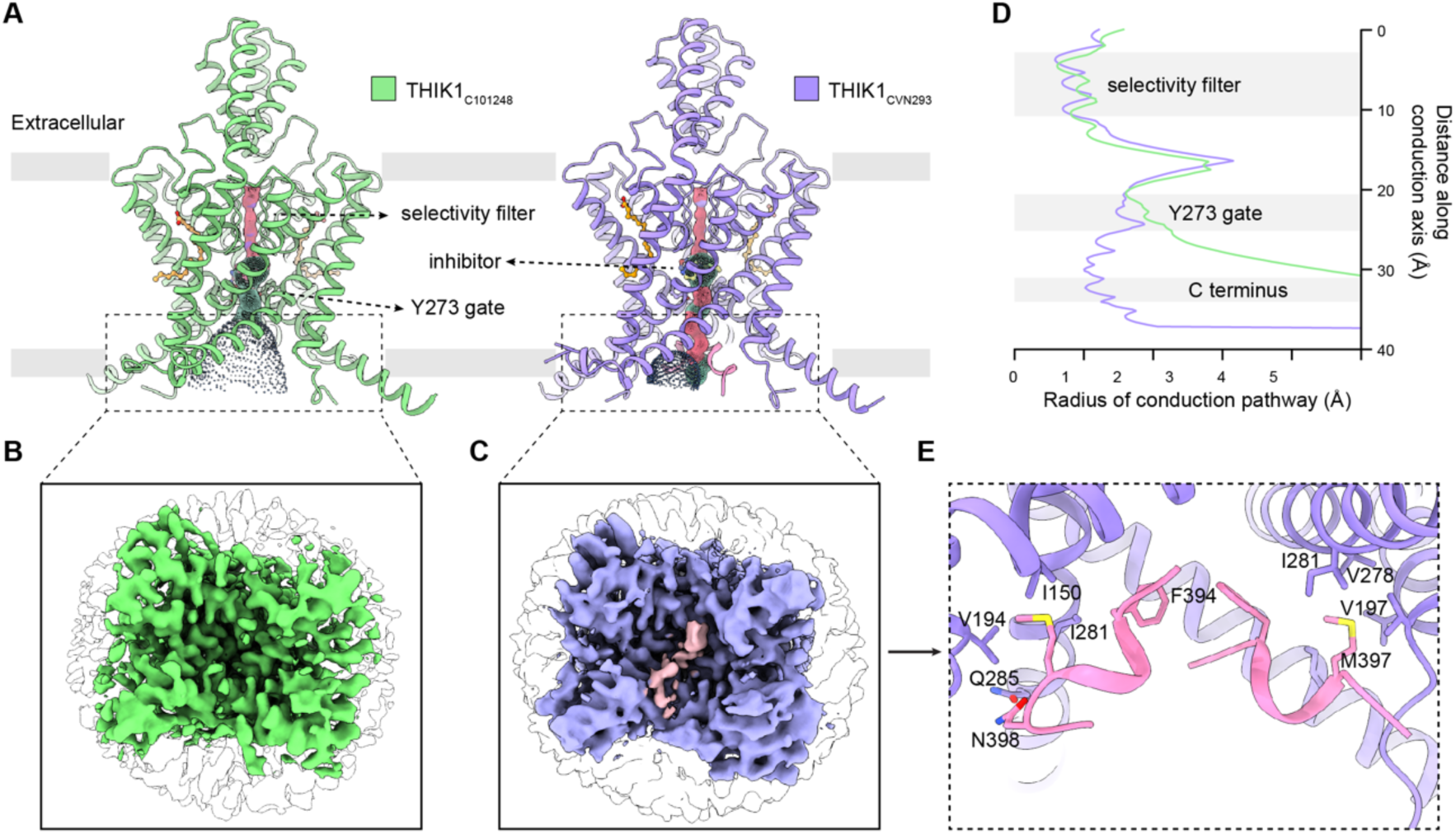
The C terminus partially resolved in THIK1_CVN293_ but absent in THIK1_C101248_. **(A)** Pore profiles of THIK1_CVN293_ and THIK1_C101248_, colored according to pore diameter. **(B-C)** Surface representations of the boxed region in **(A)**, viewed from the intracellular side, in THIK1 _C101248_ **(B)** and THIK1_CVN293_ **(C)**. The C terminus modeled in THIK1_CVN293_ is shown in pink. **(D)** Plot of pore radius for THIK1_CVN293_ (purple) and THIK1_C101248_ (light green) calculated by HOLE. **(E)** Interactions between the C terminus and adjacent residues in TM2 and TM4a in THIK1_CVN293_. An enlarged view of the C-terminal density is shown on the right.

Previous studies have shown that the C-terminal helix, located at the intracellular entrance of the vestibule, occludes K⁺ conduction and is functionally coupled to other gating elements through interactions with neighboring TM2 and TM4 (16, 27, 28). In THIK1_CVN293_, density corresponding to the C-terminal helix remained detectable, although slightly weaker than in THIK1_WT_ (Fig. 3, *A* and *C*; Fig. S6, *A* and *B*). The backbone spanning residues A393-N399 in both subunits could be modeled and was stabilized by hydrophobic interactions with nearby residues in TM2, TM4, and the TM2-TM3 loop, including I150, P192, V194, V278, and I281 (Fig. 3*E*). By contrast, the density of the C-terminal helix was markedly weaker in THIK1_C101248_, even when contoured at a lower threshold, suggesting increased conformational flexibility of the C-terminal helix upon C101248 binding (Fig. 3*B*).

In THIK1_C101248_, TM2a and TM2b undergo slight shifts toward TM1b and TM4b, respectively, accompanied by a modest upward displacement of the distal end of TM4a toward TM2b (Fig. S6*A*). Although these rearrangements are subtle, they may perturb the TM2/TM4 interface that helps anchor the C-terminal helix within the intracellular vestibule, thereby weakening its interactions with the pore and increasing its conformational flexibility. By contrast, TM2 and TM4 in THIK1_CVN293_ show minimal deviations from THIK1_WT_, preserving the interactions that stabilize the C-terminal helix at the intracellular entrance of the vestibule (Fig. 3, *A* and *C*; Fig. S6, *A* and *B*). The C-terminal helix therefore remains well defined and positioned to occlude the ion conduction pathway (Fig. 3, *C* and *D*).

Despite binding beneath the selectivity filter and interacting with surrounding residues, both inhibitors had minimal effects on the overall conformation of the selectivity filter. In both the C101248- and CVN293-bound structures, K⁺ densities remained well resolved at the S1, S3, and S4 sites, whereas the S2 site showed relatively weaker density (Fig. S7, *A* to *C*). The coordination geometry of the selectivity filter was also largely preserved, as indicated by comparable distances between the carbonyl oxygen atoms lining the filter (Fig. S7D). In addition, the lipid environment surrounding the selectivity filter was similarly preserved. Previous structural analyses have shown that linoleic acid (EIC) molecules occupying the lipid-binding pocket contribute to the stabilization of the selectivity filter (16). Consistent with this role, clear lipid densities were retained in both inhibitor-bound structures, with no apparent rearrangement of the lipid-binding environment (Fig. S7, *E* to *G*). Together, these observations indicate that the selectivity filter and its associated lipid environment remain largely unchanged upon inhibitor binding, despite the distinct effects of C101248 and CVN293 on the C-terminal gate.

### Differential roles of the C-terminal gate in inhibitor-mediated THIK1 inhibition

Beyond its roles in regulating THIK1 through intracellular signaling pathways, including caspase-8 cleavage and G protein-mediated modulation, the C terminus may also contribute to inhibitor-dependent regulation of channel activity (29, 30). As we described above, chemically related inhibitors induce distinct conformational changes at inner gate I, with the C-terminal helix remaining well defined in THIK1_CVN293_ but becoming largely unresolved in THIK1_C101248_. To exclude the possibility that the reduced C-terminal helix density in the THIK1_C101248_ structure resulted from its smaller particle number, we randomly downsampled the THIK1_CVN293_ particles to the similar number (97,657) as used for THIK1_C101248_ (97,367). The C-terminal helix remained clearly resolved in the downsampled THIK1_CVN293_ reconstruction, indicating that the weaker density in THIK1_C101248_ reflects an intrinsic conformational difference rather than lower particle count (Fig. S8). To determine whether these structural differences are functionally relevant to inhibitor action, we examined the effects of C-terminal deletion by electrophysiological recording (Fig. 4).

**Fig. 4.**
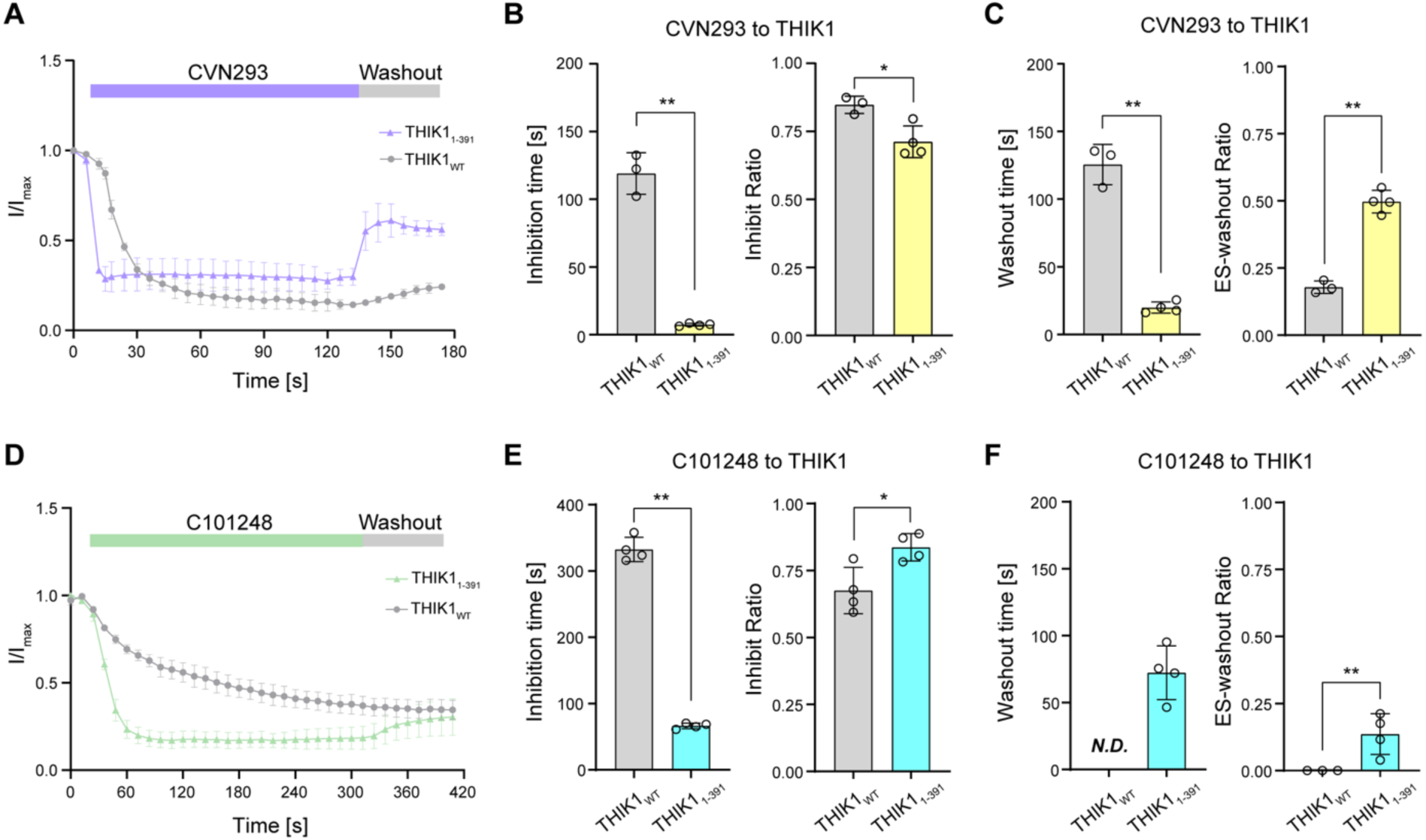
Electrophysiological responses of C-terminally truncated THIK1 to inhibitors. (A) Kinetics of CVN293 binding and dissociation in THIK1_WT_ and THIK1_1-391_. **(B)** Time required to reach the minimum current level (inhibition time) and maximal inhibition of THIK1_WT_ and THIK1_1-391_ in response to 10 μM CVN293. **(C)** Recovery kinetics and extent after extracellular solution (ES) washout in THIK1_WT_ and THIK1_1-391_. The left panel shows the time required to reach maximal recovery after washout. The right panel displays the maximal recovery extent, defined as the ratio of the ES-washout recovered current amplitude to the CVN293-inhibited current amplitude. **(D)** Kinetics of C101248 binding and dissociation in THIK1_WT_ and THIK1_1-391_. **(E)** Time required to reach the minimum current level (inhibition time) and maximal inhibition of THIK1_WT_ and THIK1_1-391_ in response to 1 μM C101248. **(F)** Recovery kinetics and extent after extracellular solution (ES) washout in THIK1_WT_ and THIK1_1-391_. The left panel shows the time required to reach maximal recovery after washout. The right panel displays the maximal recovery extent, defined as the ratio of the ES-washout recovered current amplitude to the C101248-inhibited current amplitude. N.D., not detected. *P < 0.05 or **P < 0.01 compared with THIK1_WT_, unpaired Student’s t-test. Data are presented as the mean ± SD, with individual symbols represent individual cells tested in each group.

**Fig. 5.**
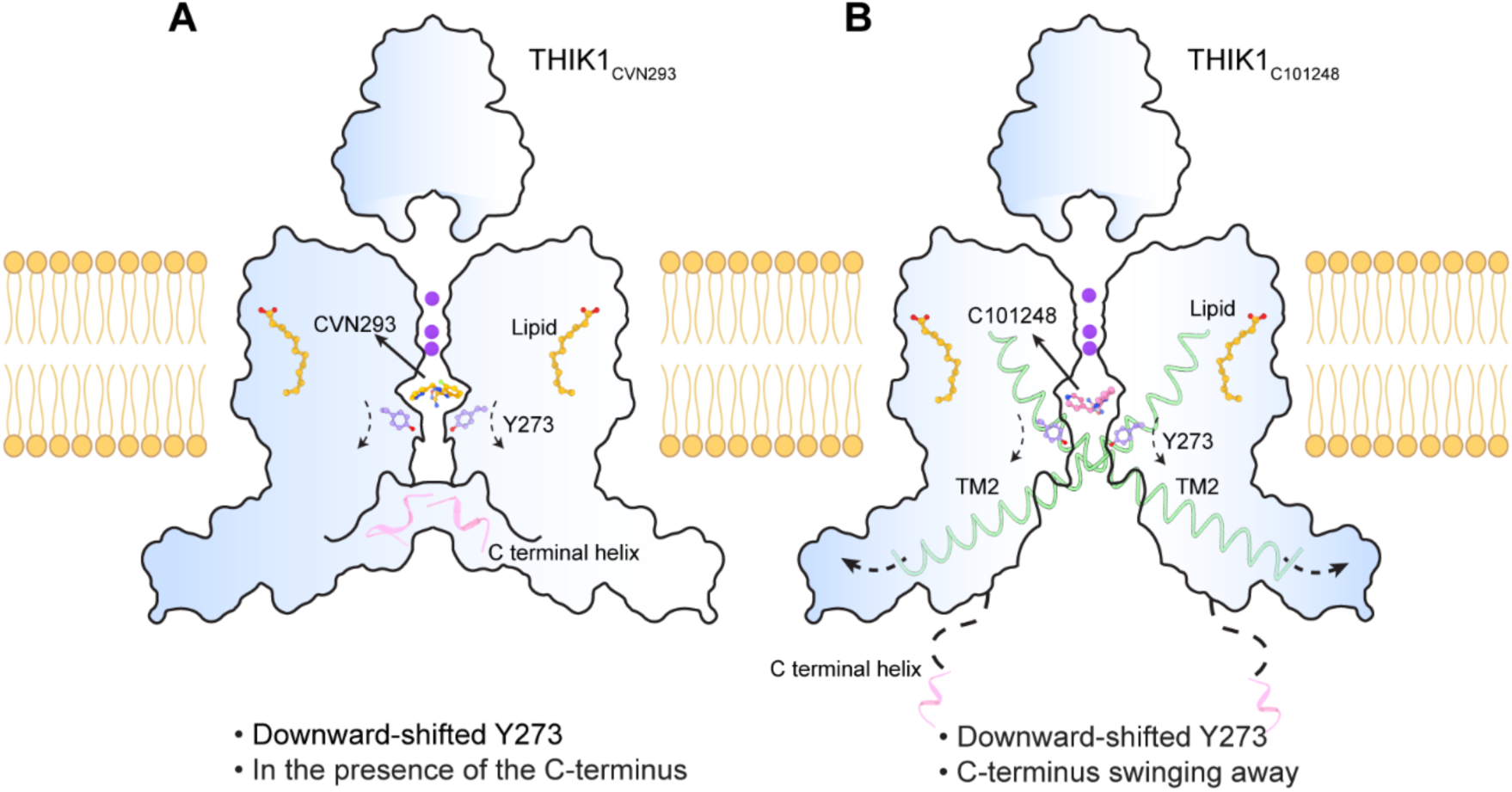
Schematic model of THIK1 inhibition by CVN293 and C101248. **(A)** Schematic model of CVN293-bound THIK1. K^+^ ions and lipid molecules are stably accommodated within the channel, Y273 undergoes a downward shift, and the C-terminal inner gate adopts a closed conformation. **(B)** Schematic model of C101248-bound THIK1. K^+^ ions and lipid molecules are stably accommodated within the channel, Y273 undergoes a downward shift, and the C-terminal helix–formed inner gate becomes flexible. K^+^ ions are colored purple, lipid molecules orange, Y273 light purple, the C-terminal helix pink, and TM2 of THIK1_C101248_ light green.

Removal of the C-terminal helix (G392-R408), a key structural component of inner gate I, reduced the maximal inhibition mediated by CVN293 from 84.8 ± 3.2% in THIK1_WT_ to 71.2 ± 5.9% in THIK1_1-391_(Fig. 4, A and B, Fig. S9). Notably, THIK1_1–391_ exhibited markedly faster inhibition kinetics, reaching maximal inhibition within 7.36 ± 1.00 s compared with 119 ± 15 s for THIK1_WT_ (Fig. 4, *A* and *B*). These findings suggest that removal of the C-terminal helix facilitates access of CVN293 to its inhibitory site. Consistent with this interpretation, CVN293 dissociated substantially faster from THIK1_1–391_, as reflected by the shorter washout time and more complete recovery of channel activity following inhibitor removal (Fig. 4, *A* and *C*). Thus, although the C-terminal helix is not required for CVN293 inhibition, its presence appears to stabilize the inhibited state and slow inhibitor dissociation. Together with the previously reported role of Y273 at inner gate II in inhibitor action, our findings identify inner gate I as an additional structural element that modulates the functional response of THIK1 to inhibitor binding (23).

Deletion of the C-terminal helix also accelerated the inhibitory response to C101248, with THIK1_1-391_ reaching its minimum current level substantially faster than THIK1_WT_ (Fig. 4, *D* and *E,* Fig. S9). Notably, while C101248-mediated inhibition was completely resistant to washout in THIK1_WT_, deletion of the C-terminal helix permitted only a marginal degree of recovery, with merely a small fraction of the current being reversed upon washout (Fig. 4*F*). This indicates that the C-terminal helix does influence C101248 dissociation, but its contribution is rather limited. In contrast to C101248, CVN293 inhibition, which was relatively resistant to washout in THIK1_WT_, became effectively reversible upon C-terminal deletion, highlighting distinct inhibitor-specific dependencies on the C-terminal gate. The modest effect of the C-terminal helix on C101248 inhibition and dissociation is consistent with the structural observation that this helix is poorly resolved in the THIK1_C101248_ complex, implying that the helix is not critically required for stabilizing C101248 binding or inhibitory efficacy. Rather than requiring a defined open conformation of inner gate I, these findings support a model in which the conformational flexibility of the C-terminal gate modulates inhibitor access and efficacy in an inhibitor-dependent manner.

Overall, C101248- and CVN293-mediated inhibition of THIK1 is differentially modulated by the integrity of inner gate I. An intact inner gate I enhances the extent and persistence of CVN293-mediated inhibition, as reflected by stronger inhibition and slower inhibitor dissociation. In contrast, removal of the C-terminal helix enhances C101248-mediated inhibition, consistent with the increased flexibility of this gate observed in the C101248-bound structure. These findings indicate that, despite their close chemical relationship and shared binding site, C101248 and CVN293 can be differentially influenced by the conformational state of the C-terminal gate, highlighting the ability of a common regulatory element to distinguish between chemically related inhibitors.

## Discussion

THIK1 belongs to the K2P family and is critical for maintaining microglial membrane potential and potassium homeostasis (2, 31). Its activity is regulated by diverse intracellular and extracellular cues, including G protein-coupled receptor pathways, caspase-8-mediated cleavage, and lipid interactions (29, 30, 32). In response to regulatory inputs, the inner vestibule, selectivity filter, and intracellular C terminus engage in coordinated structural rearrangements that together define an integrated gating network (16, 23, 33).

Here, we investigated how two chemically related inhibitors, C101248 and CVN293, inhibit THIK1. Although the two compounds share a related chemical scaffold and bind within the same inner vestibular region beneath the selectivity filter, their effects on the channel are not identical. Both inhibitors are associated with remodeling of the Y273 inner gate (inner gate II), whereas the selectivity filter and its associated lipid environment remain largely preserved. The most prominent difference between the two inhibitor-bound structures is observed at the C-terminal helix that forms inner gate I. The C-terminal helix remains well defined in the CVN293-bound structure but becomes poorly resolved in the C101248-bound structure, consistent with increased conformational flexibility upon C101248 binding. The particles in THIK1 These observations indicate that the two inhibitors produce distinct conformational responses despite engaging a common binding site.

Our electrophysiological analyses further support a functional role for the C-terminal gate in inhibitor-mediated inhibition. Removal of the C-terminal helix accelerated the inhibitory response to both compounds, suggesting that the integrity of this gate influences access to or engagement of the inhibitory site. However, the consequences of C-terminal deletion differed between the two inhibitors. For CVN293, deletion modestly reduced the extent of inhibition and substantially accelerated washout, indicating that the C-terminal helix contributes to maintaining the inhibited state. In contrast, C-terminal deletion enhanced the extent of C101248-mediated inhibition, consistent with the increased flexibility of this region observed in the C101248-bound structure. Thus, the C-terminal gate does not simply promote or oppose inhibitor action; rather, its functional contribution depends on the inhibitor.

Thus, our findings show that chemically related inhibitors sharing a common binding site can nevertheless engage the same regulatory element differently. This difference is consistent with the distinct conformations of the C-terminal helix observed in the two inhibitor-bound structures and suggests that subtle differences in ligand chemistry can influence the conformation of a shared regulatory element. Together with the previously described role of the Y273 inner gate (23), our findings suggest the inner vestibule as an important region for inhibitor-mediated regulation of THIK1. Notably, the selectivity filter and its associated lipid environment remain largely unchanged in both inhibitor-bound structures, indicating that the C-terminal gate can respond differently to inhibitor binding without major structural rearrangement of the selectivity filter. Overall, these findings provide structural and functional insights into how chemically related inhibitors can modulate THIK1 differently despite engaging a common binding site.

## Materials and Methods

### Protein expression and purification of THIK1

The full-length Homo sapiens THIK1 cDNA was amplified from a human cDNA library and cloned into the modified pBMCL1 vector (34, 35), followed by a PreScission protease cleavage site, green fluorescent protein (GFP) and a Strep tag at the C terminus.

THIK1 was expressed in 1L HEK293S GnTI⁻ cells and cultured in FreeStyle™ 293 medium (Thermo Fisher Scientific) at 37°C with shaking at 110 rpm under 8% CO_2_. When the cell density reached around 2.5×10^6^ cells/ml, 1 mg of plasmid DNA encoding recombinant THIK1 and 3 mg of PEI transfection reagent (PolyScience) were mixed with 50 mL of culture medium. After incubation at room temperature for 30 min, the mixture was added to the culture. 16 h later, sodium butyrate was added to a final concentration of 10 mM, and the temperature was reduced to 30°C. Cells were harvested after another 48 h by centrifugation at 2,000g for 10 min, washed once with ice-cold PBS, then immediately used for protein purification.

The harvested cells were resuspended in 50 ml lysis buffer (20 mM Tris, pH 7.8, 150 mM KCl, 2 mM DTT, 1% DDM, 0.2% CHS, protease inhibitors, and 2 μg/ml DNase I). The mixture was homogenized using a glass Douncer for 20 min and then gently stirred at 4°C for 2 h. After pelleting cell debris at 50,000g for 45 min, the supernatant was collected and incubated with 1.5 ml of Streptactin beads 4FF (Smart-Lifesciences) for 2 h at 4°C. The resin was loaded onto a gravity column and washed with 100 ml wash buffer I (20 mM Tris, pH 7.8, 150 mM KCl, 2 mM DTT, 0.025% DDM, 0.005% CHS). Prior to elution, the resin was briefly equilibrated in wash buffer II (20 mM Tris, pH 7.8, 150 mM KCl, 2 mM DTT, 0.01% LMNG, 0.001% CHS) at 4°C for 20min. The protein was eluted with 15ml wash buffer II containing 7.5 mM D-desthiobiotin. The elution was concentrated to 0.7ml and centrifuged at 13,000 rpm for 10 min. The supernatant was further purified by size-exclusion chromatography (SEC) on a Superose 6 increase 10/300 GL column (Cytiva) pre-equilibrated with SEC buffer (20 mM Tris, pH 7.8, 150 mM KCl, 2 mM DTT, 0.01% LMNG, 0.001% CHS). The peak fraction was collected and concentrated to 2 mg/mL for cryo-EM.

### Cryo-EM sample preparation

The protein sample was centrifuged at 17,000g for 10 min at 4 °C. CVN293 and C101248 were added separately to the protein samples at final concentrations of 750 μM and 100 μM, respectively, followed by incubation on ice for 1 h prior to grid preparation. 3 µl of the protein sample was applied to holey carbon, 300-mesh R1.2/1.3 gold grids (Quantifoil) that were freshly glow-discharged for 30 s. Cryo-EM grids were prepared on a Vitrobot Mark IV (Thermo Fisher Scientific) under controlled conditions of 4 °C and 100% humidity. After 5 s incubation, the grids were blotted for 3 s with filter paper and then rapidly plunge-frozen in liquid ethane cooled by liquid nitrogen.

### Cryo-EM data acquisition and processing

Cryo-EM grids were loaded into a Titan Krios G4 microscope (Thermo Fisher Scientific) operated at 300 kV and equipped with a Falcon G4i direct electron detector. Data were acquired automatically using EPU software in super-resolution mode. Movies were recorded with a total exposure time of 3.51 s with a calibrated pixel size of 0.932 Å. Each EER stack was collected with an accumulated dose of ∼50 e^-^Å^-2^.

For the THIK1_CVN293_ dataset, 4,229 micrographs were collected. Each EER movie stack, consisting of 1,080 frames, was partitioned into 27 fractions and corrected for beam-induced drift with MotionCor2 implemented in RELION (version 4.0.0) (36, 37). Following upsampling and binning, the images were processed to a pixel size of 0.932 Å per pixel. The exposure-weighted micrographs were subsequently imported into cryoSPARC (version 4.5.3) (38), and CTF estimation was performed using Patch CTF. Blob picker was conducted on 50 micrographs, using a circular diameter of 80-160 Å. After initial 2D classification, high-quality templates were selected for subsequent template picker. Several rounds of 2D classification on all micrographs were carried out to refine particle selection. Given the limited representation of side-view particles in the particle set, Topaz was trained using manually selected particles to enrich the proportion of side-view particles during subsequent particle picking. A total of 518,691 particles were subjected to ab initio reconstruction with three classes, with no symmetry applied. To improve the resolution, particles belonging to the best-defined class underwent another round of ab initio reconstruction and non-uniform refinement. A final resolution of 2.81 Å was determined from the gold-standard Fourier shell correlation (FSC) using the 0.143 cutoff, and the map was further sharpened by applying a B-factor of -60.

For the THIK1_C101248_ dataset, EER movie stacks of 3,524 frames were acquired and processed in a manner analogous to the THIK1_CVN293_ dataset, achieving a final map with an overall resolution of 2.99 Å.

### Model building, refinement, and validation

De novo atomic model building of THIK1 was carried out in Coot (39) based on the reconstructed cryo-EM maps. Atomic model building of THIK1 was initiated using the full-length THIK1 (PDB ID: 9JGZ) structure as a starting template. Real-space refinement was carried out using phenix.real_space_refine against the cryo-EM density maps, with Ramachandran and non-crystallographic symmetry restraints applied (40). Model geometry and stereochemical quality were evaluated using MolProbity (41). Pore dimensions were calculated with the HOLE program (42). Protein–ligand interactions were analyzed using LIGPLOT (43), and all structural figures were generated in ChimeraX (44).

### Whole-cell patch-clamp electrophysiology

Human embryonic kidney (HEK293) cells were maintained in Dulbecco’s Modified Eagle Medium (DMEM; Gibco, USA) containing 1% Glutamax (Gibco, USA), 1% penicillin‒ streptomycin (Gibco, USA), and 10% fetal bovine serum (FBS; WISENT, Canada). Cell culture conditions were set at 37 °C and a humidified atmosphere of 5% CO_2_. For plasmid delivery, the calcium phosphate transfection method was utilized. Patch pipettes were fabricated from glass capillaries using a two-stage vertical puller (PC-100, Narishige, Japan) to yield a tip resistance ranging from 3 to 5 MΩ when loaded with the internal solution. This pipette solution contained (in mM) 120 KCl, 30 NaCl, 0.5 CaCl_2_, 1 MgCl_2_, 10 HEPES, and 5 EGTA (pH 7.2). The extracellular solution (ES) was composed of (in mM) 150 NaCl, 5 KCl, 10 glucose, 2 CaCl_2_, 10 HEPES, and 1 MgCl_2_ (pH 7.35-7.40). Whole-cell recordings were carried out at room temperature (25 ± 2 °C) within 24-48 h post-transfection. All currents were sampled using an Axopatch 200B amplifier coupled with a Digidata 1550B digitizer (Molecular Devices, USA), digitized at 10 kHz and low-pass filtered at 2 kHz. Data were acquired and analyzed using Clampex 10.7 (Molecular Devices, USA) and prism 10. C101248 or CVN293, prepared in the extracellular solution, was applied through a Y-tube.

## Acknowledgments

We thank the Cryo-Electron Microscopy Facility of Henan University and the Cryo-EM Center at Fudan University, as well as the staff at both facilities, for their technical assistance.

## Fundings

Brain Science and Brain-like Intelligence Technology-National Science and Technology

Major Project 2022ZD0207800 (B.L., J.W.)

National Natural Science Foundation of China grant 32371261 and 32671280 (B.L.)

National Natural Science Foundation of China grant 32301011 (R.Z.)

National Natural Science Foundation of China grant 32471203 and 32671536 (J.W.)

Fundamental Research Funds for the Central Universities grant 2632026TD02 (J.W.)

Natural Science Foundation of Hangzhou for Young Scholars grant 2025SZRJJ1838 (J.W.)

## Author Contributions

B.L., J.W. and R.Z. (Ran Zhang) conceived and supervised the project. X.F. prepared the samples; R.Z. (Ruiheng Zhang) and X.F. performed data acquisition, image processing and structure determination; R-H.Z., H.J., B.W. and J.W. performed electrophysiology and analyzed the data; X.F., Z.W., W.W. and X.Z. contributed to the methodology. B.L. and X.F. wrote the manuscript with the input from all authors.

## Supplementary Materials

**Fig. S1.**
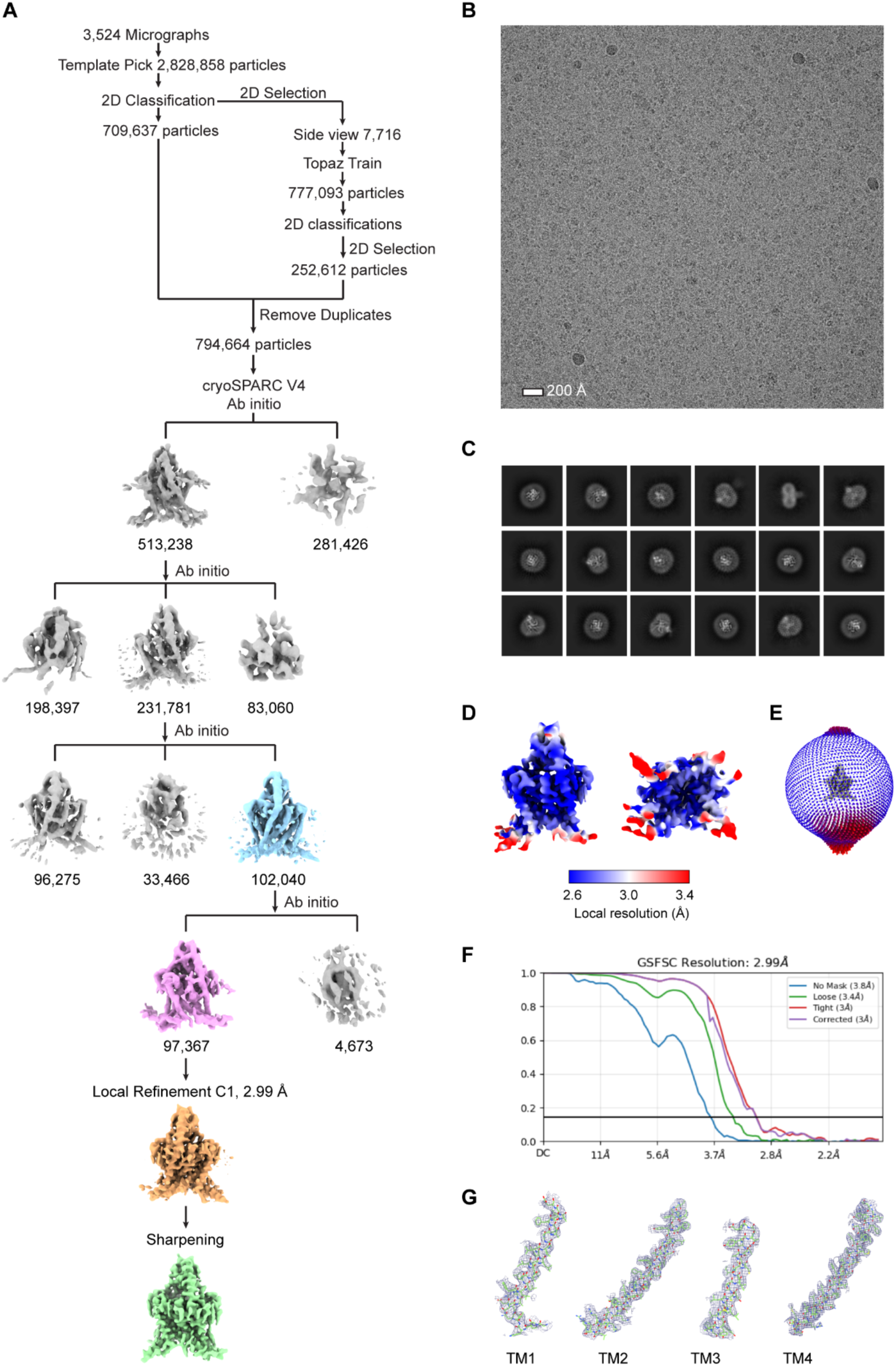
Cryo-EM processing pipeline for THIK1_C101248._ **(A)** Cryo-EM processing workflow for THIK1_C101248_ in cryoSPARC4.4.1. See Methods for details. **(B)** Representative micrograph of THIK1_C101248_. **(C)** 2D class average of THIK1_C101248_ particles. **(D)** Local resolution distribution of THIK1_C101248_ map. **(E)** Angular distribution of particles used in final reconstruction. **(F)** The GSFSC curve of the final map. **(G)** Cryo-EM densities for transmembrane helices in THIK1_C101248_.

**Fig. S2.**
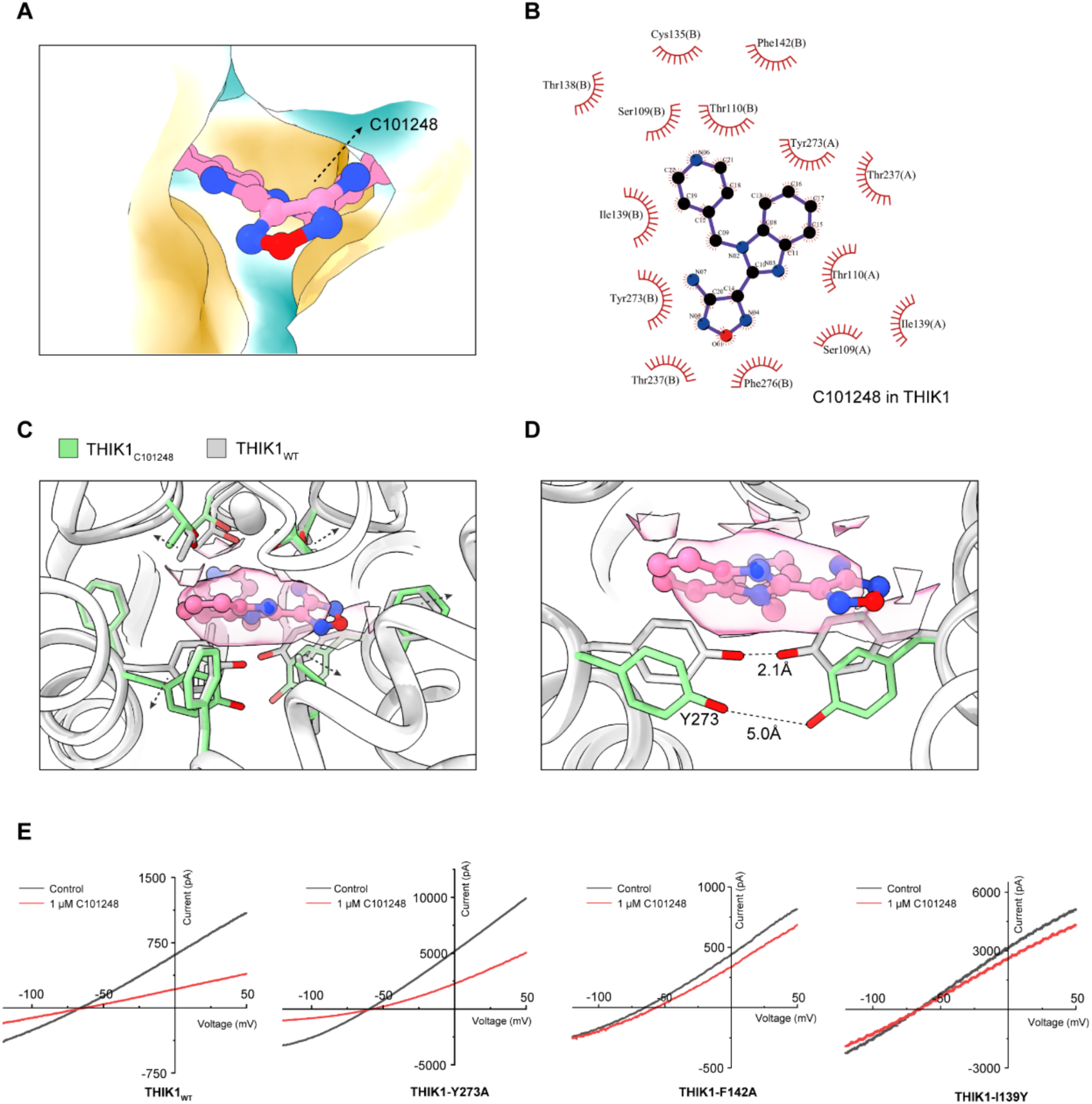
C101248 occupancy in the inner vestibule. **(A)** Hydrophobic surface representation of the C101248 binding pocket (hydrophilic regions colored blue, transitioning to white and then to orange for the most hydrophobic regions). **(B)** Ligplot analysis of C101248 in the modulator pocket, with key residues labeled. **(C-D)** Residues forming key interactions with C101248 exhibit a subtle outward shift **(C)**, particularly Y273 displaying the most pronounced downward displacement **(D)**. Ribbon model of THIK1_C101248_ is colored in white and its stick is shown in light green. Model of THIK1_WT_ is colored in gray. C101248 is shown in pink, with its corresponding density displayed as a transparent surface. **(E)** Representative inhibitory current traces of different mutants.

**Fig. S3.**
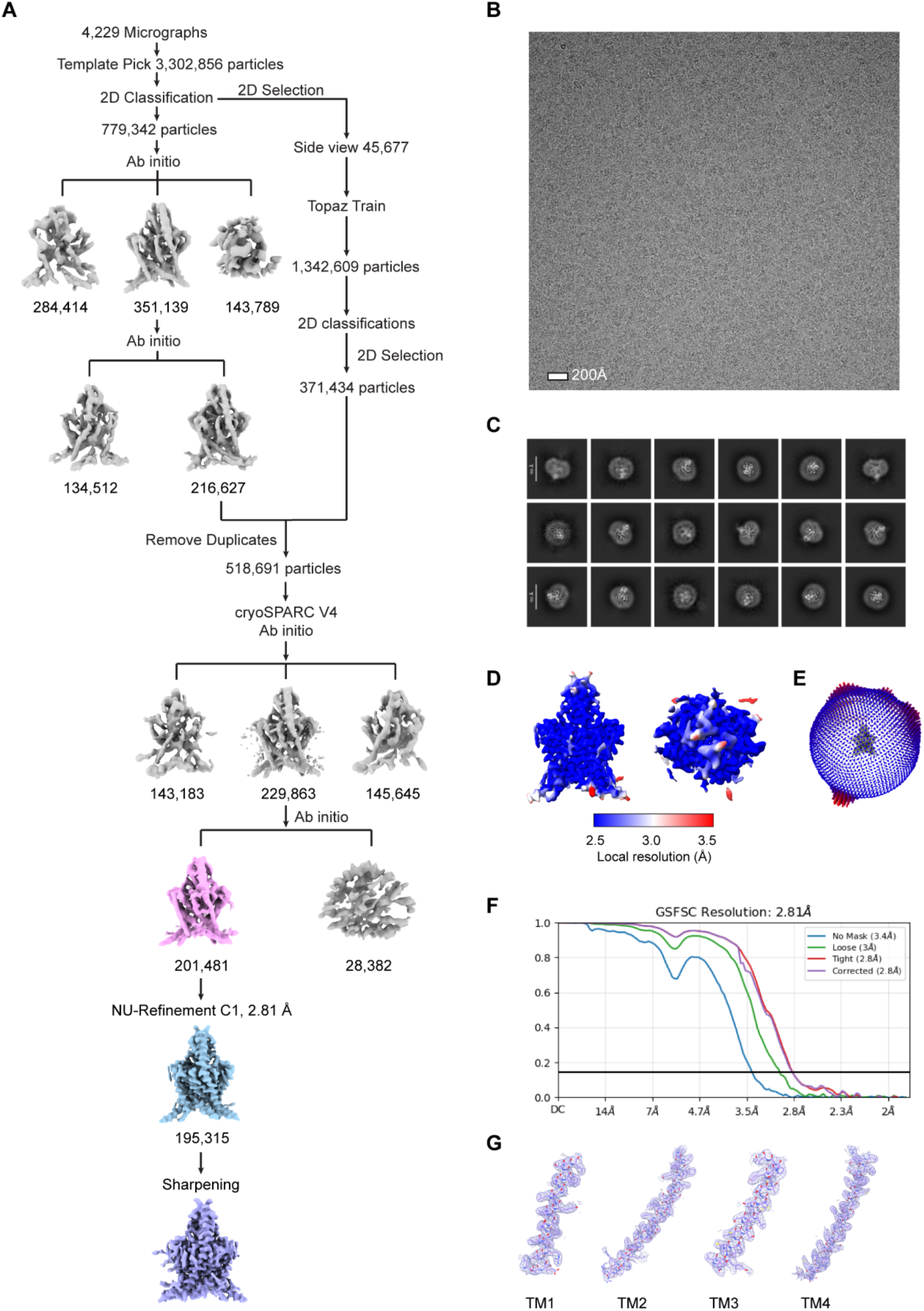
Cryo-EM processing pipeline for THIK1_CVN293._ **(A)** Cryo-EM processing scheme for THIK1_CVN293_ in cryoSPARC4.4.1. See Methods for details. **(B)** Representative micrograph of THIK1_CVN293_. **(C)** Representative 2D class average particles of THIK1_CVN293_. **(D)** Local resolution distribution of THIK1_CVN293_ map. **(E)** Angular distribution of particles for the final reconstruction. **(F)** The GSFSC curve of the final map. **(G)** Representative densities for THIK1_CVN293_ transmembrane helices.

**Fig. S4.**
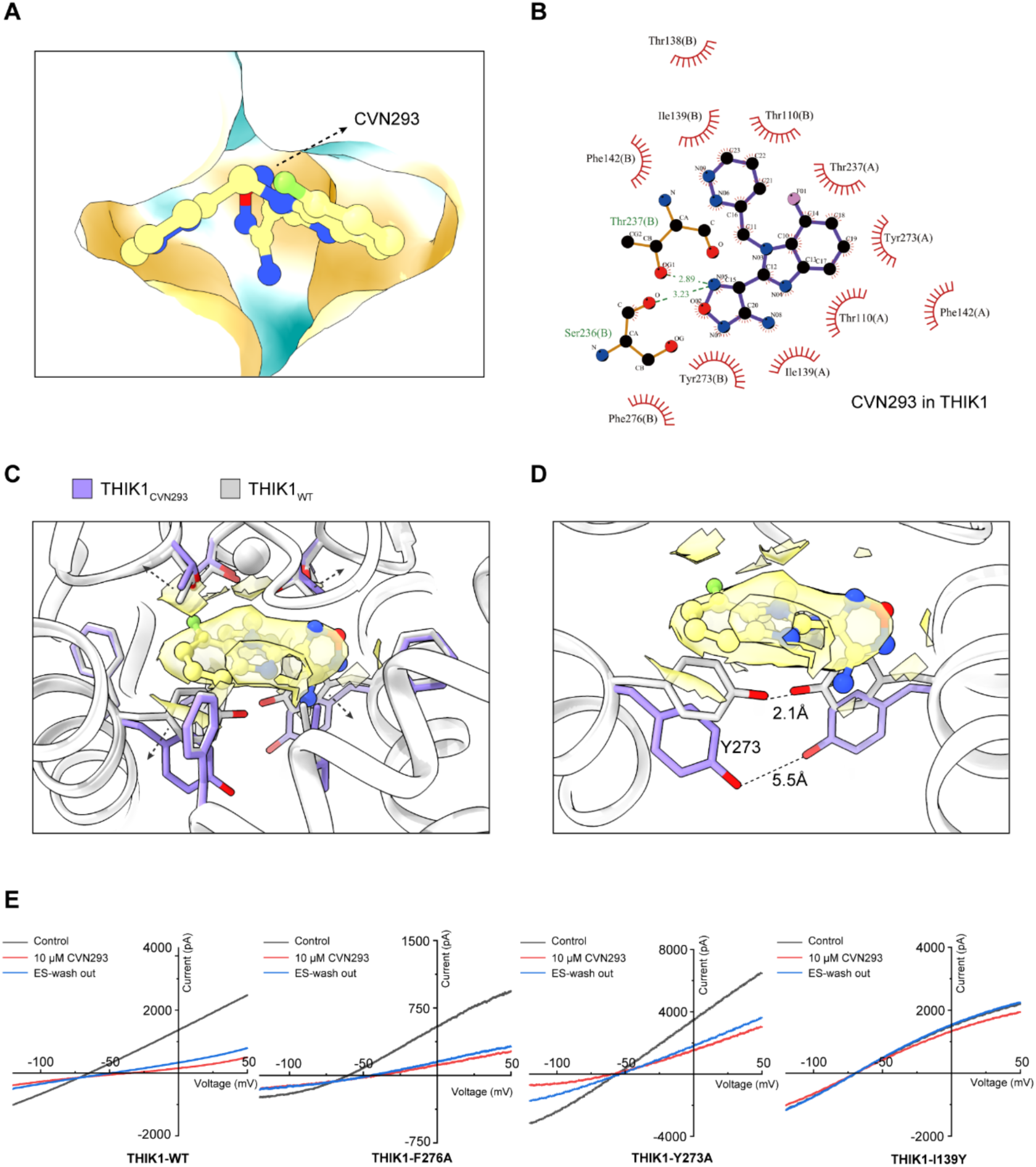
CVN293 occupancy in the inner vestibule. **(A)** Hydrophobic surface representation of the CVN293 binding pocket (hydrophilic regions colored blue, transitioning to white and then to orange for the most hydrophobic regions). **(B)** Ligplot analysis of CVN293 in the modulator pocket, with key residues labeled. **(C-D)** Residues forming key interactions with CVN293 exhibit a subtle outward shift **(C)**, particularly Y273 displaying the most pronounced downward displacement **(D)**. Ribbon model of THIK1_CVN293_ is colored in white and its stick is shown in purple. Model of THIK1_WT_ is colored in gray. CVN293 is shown in pink, with its corresponding density displayed as a transparent surface. **(E)** Representative inhibitory current traces of different mutants.

**Fig. S5.**
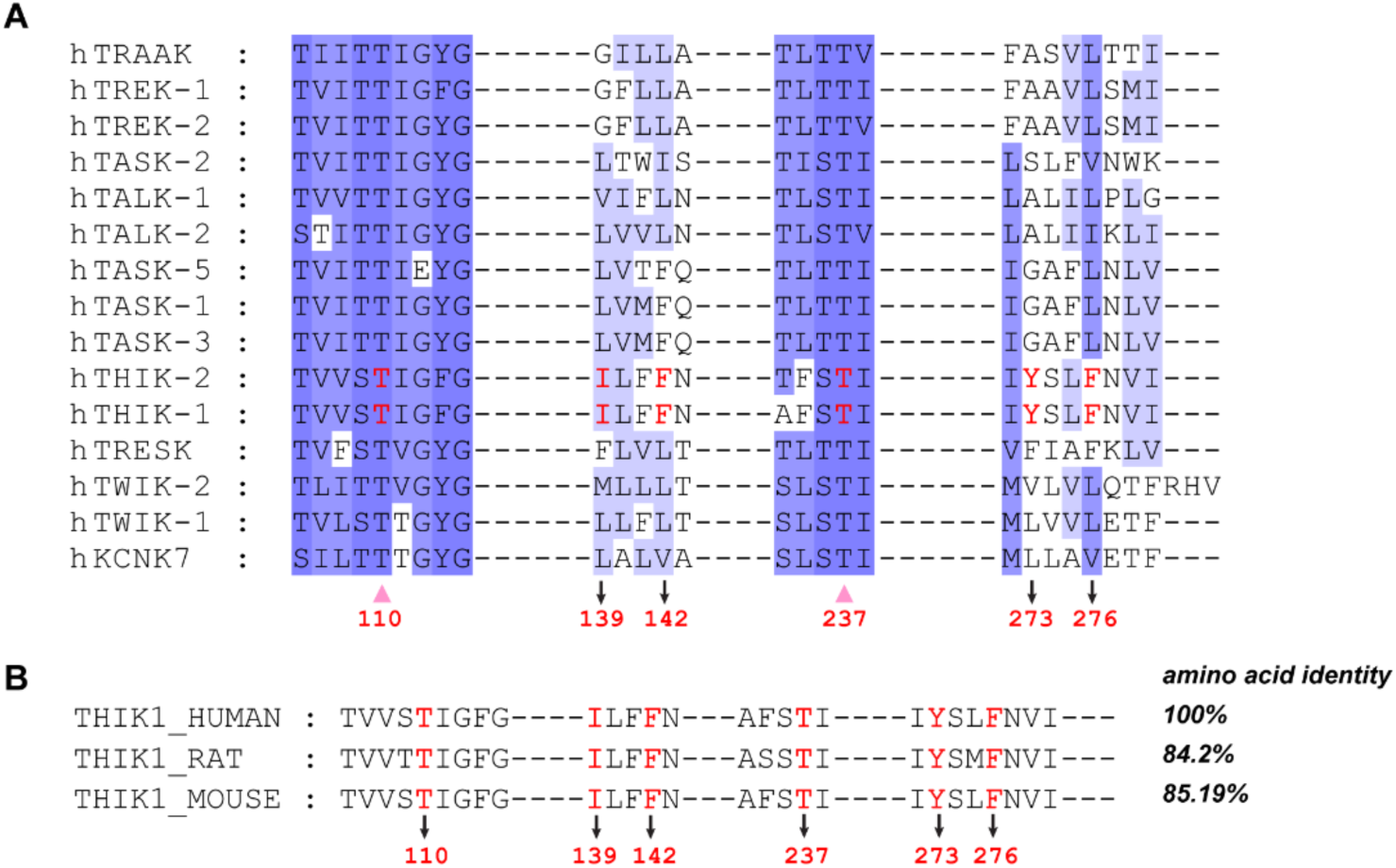
Conservation analysis of THIK1 residues commonly engaged by C101248 and CVN293. **(A)** Sequence conservation of the binding pocket shared by C101248 and CVN293 across the K2P family, with conserved residues highlighted in bold red. The residue column marked by the bottom pink triangle is located in the SF and highly conserved in K2P family. **(B)** Sequence alignment of human, rat and mouse THIK1 orthologues, showing high overall sequence similarity and strong conservation of inhibitor-binding residues.

**Fig. S6.**
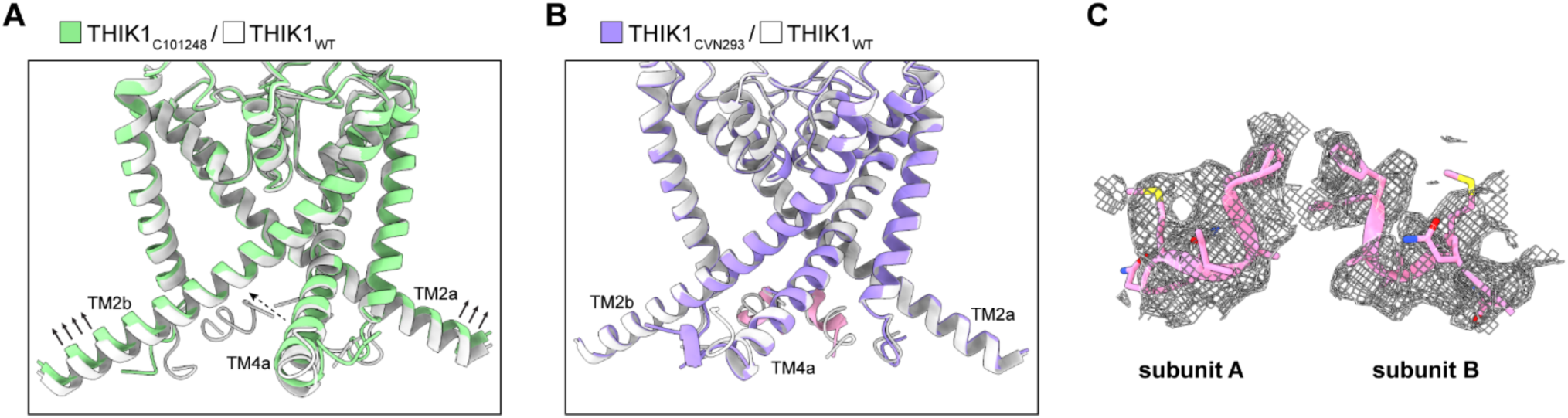
C101248-induced conformational rearrangements in TM2 and TM4 increase C-terminal flexibility. **(A)** Relative to inhibitor-free THIK1, the distal segments of TM2 and TM4a lift upward in THIK1_C101248_. TM2 and TM4a in THIK1_CVN293_ retain conformations closely resembling those observed in inhibitor-free THIK1. **(B)** An enlarged view of the C-terminal density is shown on the right. **(C)** Densities of C -terminal helix of THIK1CVN293.

**Fig. S7.**
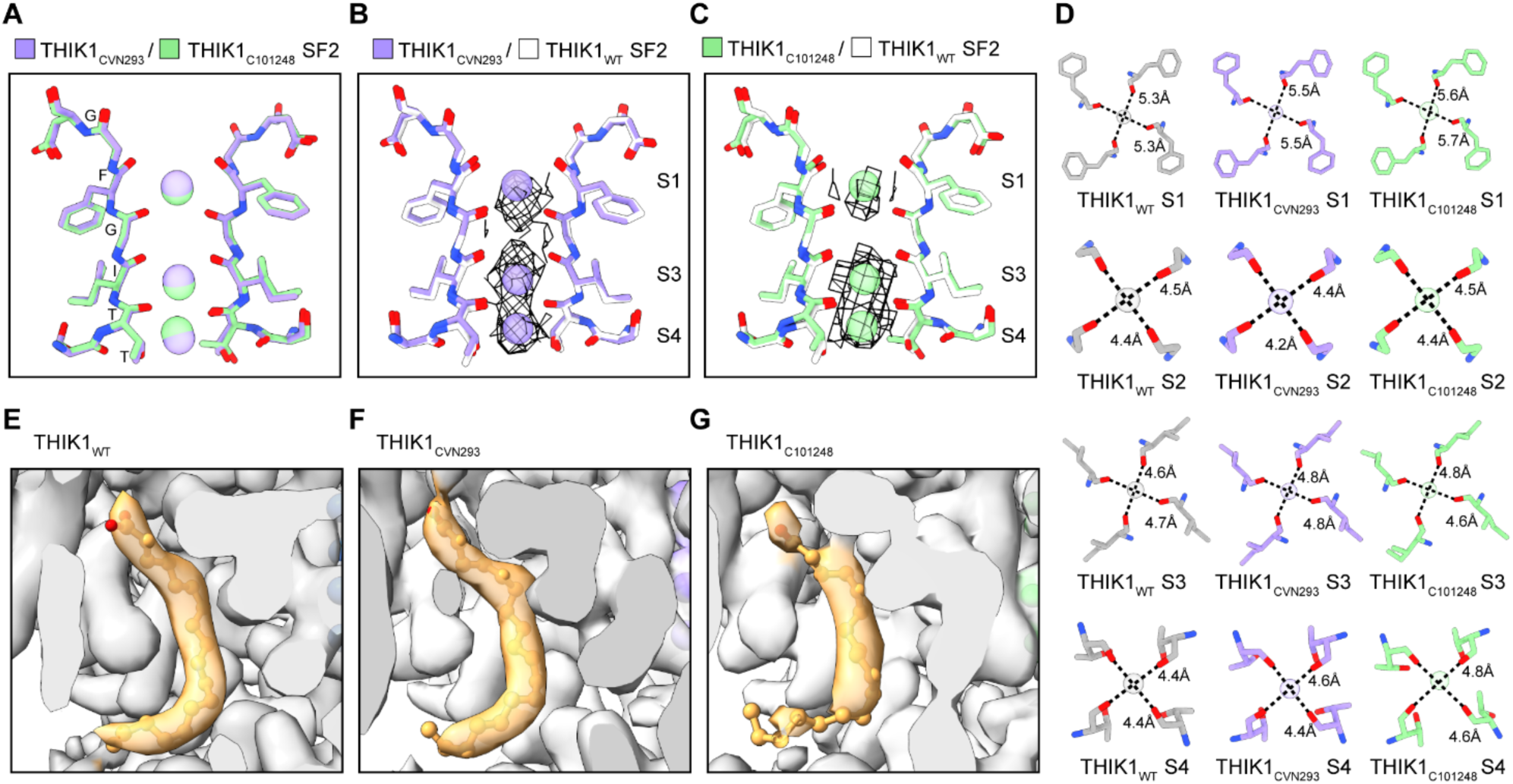
Distinct effects of CVN293 and C101248 on selectivity filter conformation and lipid binding in THIK1. **(A)** Superposition of the SF2 region from THIK1_CVN293_ and THIK1_C101248_. **(B)** Superposition of the SF2 region of THIK1_CVN293_ with THIK1_WT_ (white). K⁺ ion densities are shown. **(C)** Superposition of the SF2 region of THIK1_C101248_ with THIK1_WT_. K⁺ ion densities are shown. **(D)** Inter-carbonyl distances measured at the S1, S2, S3 and S4 K⁺-binding sites in THIK1_WT_, THIK1_CVN293_ and THIK1_C101248_. **(E-G)** Density corresponding to the bound lipid in THIK1_WT_, THIK1_CVN293_ and THIK1_C101248_.

**Fig. S8.**
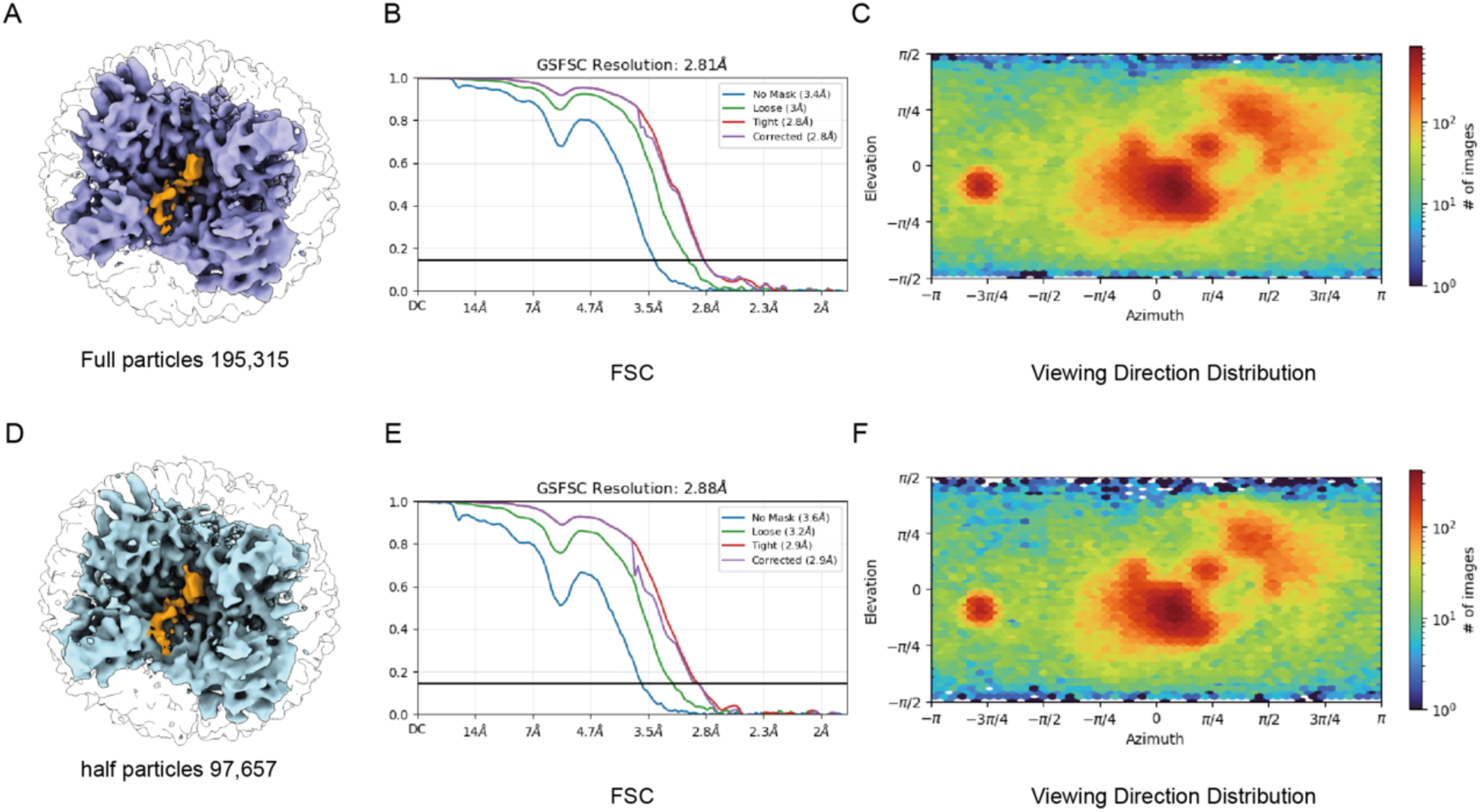
The C-terminal helix of THIK1_CVN293_ remains resolved after particle downsampling. (A, D) Surface representations of THIK1_CVN293_ viewed from the intracellular side, reconstructed from the full particle set (A, 195,315 particles) or the randomly downsampled half particle set (D, 97,657 particles). The C-terminal helix remains well resolved in the downsampled map. (B, E) GSFSC curves of the corresponding final maps. (C, F) Euler angle distributions of particles for the two reconstructions.

**Fig. S9.**
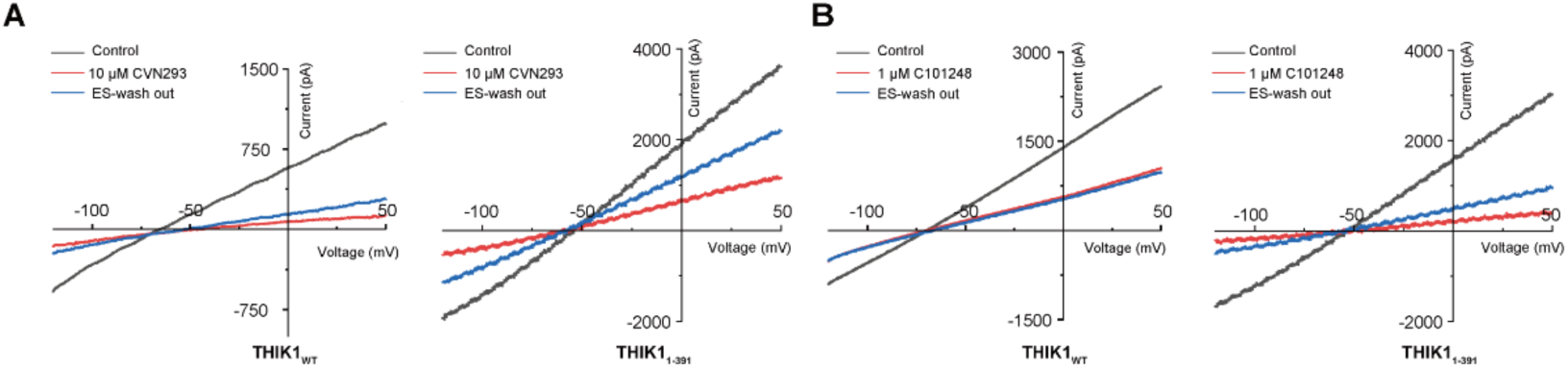
Electrophysiological analysis of CVN293 and C101248 inhibition of THIK1. **(A and B)** Current-voltage (I-V) curves during CVN293 **(A)** or C101248 **(B)** Binding and washout. Baseline traces are shown in grey, inhibition traces in red, and washout traces in blue.

